# Analysis of antennal responses to motion stimuli in the honey bee by automated tracking using DeepLabCut

**DOI:** 10.1101/2023.04.24.538069

**Authors:** Hiroki Kohno, Shuichi Kamata, Takeo Kubo

## Abstract

Considering recent developments in gene manipulation methods for honey bees, establishing simple, robust, and indoor assay systems which can analyze behavioral components in detail is important for the rise of honey bee behavioral genetics. We focused on the movements of antennae of the honey bee, which are used for not only multimodal sensory perception but also interactions between individuals. We developed an experimental system for analyzing the antennal responses (ARs) of the honey bee using DeepLabCut, a markerless posture-tracking tool using deep learning. The tracking of antennal movements during the presentation of vertical (downward and upward) motion stimuli using DeepLabCut successfully detected the ARs reported in the previous studies, where bees tilted their antennae in the direction opposite to the motion stimuli. In addition, we successfully detected ARs in response to horizontal (forward and backward) motion stimuli. An investigation of the developmental maturation of honey bee ARs showed that ARs to motion stimuli were not detected in bees immediately after emergence but became detectable through post-emergence development in an experience-independent manner. Furthermore, unsupervised clustering analysis using multidimensional data created by processing tracking data using DeepLabCut classified antennal movements into different clusters, suggesting that data-driven behavioral classification can apply to AR paradigms. These results reveal novel AR to visual stimuli and developmental maturation of ARs and suggest the efficacy of data-driven analysis for behavioral classification in behavioral studies of the honey bee.

**Summary statement:** Automated tracking using DeepLabCut was successfully applied to measure the antennal response to motion stimuli and unsupervised classification of antennal movements in honey bees.

## Introduction

Genome sequencing and comprehensive gene expression analyses have been conducted on many insect species (Ellegren, 2014; Oppenheim et al., 2015). Combined with the establishment of gene manipulation methods (Adli, 2018; Mello and Conte, 2004), the molecular and neural bases of insect behavior have been elucidated in various non-model insect species other than *Drosophila* (Mansourian et al., 2019; Sun et al., 2017; Walton et al., 2020). The European honey bee (*Apis mellifera* L.) is a well-known social insect, and its social behavior has been extensively studied for many years (Frisch et al., 1967; Seeley, 1995; Winston, 1987). In addition, genome editing and transgenic technologies have been established and applied for molecular- and neuro-ethological analyses in honey bees (Carcaud et al., 2023; Kohno and Kubo, 2018; Kohno and Kubo, 2019; Kohno et al., 2016; Otte et al., 2018; Roth et al., 2019; Schulte et al., 2014), although the molecular and neural basis underlying honey bee behaviors remain largely unknown.

Most innate behaviors of honey bees, such as nursing their brood, division of labor of workers, and waggle dance, have been described by observations and behavioral experiments in the field (Frisch et al., 1967; Seeley, 1995), but genetically modified honey bees must be confined to laboratory conditions due to legal restrictions. To date, a simple and robust behavioral experimental paradigm, olfactory conditioning of the proboscis extension reflex (PER), has been extensively utilized for the indoor analysis of the abilities and mechanisms of learning and memory in honey bees (Eisenhardt, 2014; Giurfa and Sandoz, 2012). Some previous studies have developed original devices for analyzing bee behaviors and sophisticated psychological experimental paradigms (Geng et al., 2022; Giurfa et al., 2001; Howard et al., 2018; Howard et al., 2019; Kirkerud et al., 2013; Kirkerud et al., 2017; Marchal et al., 2019; Nouvian and Galizia, 2019; Schultheiss et al., 2017), but they have not necessarily been widely used due to their uniqueness or difficulties requiring skilled handling of bees with care. Against this background, developing a variety of robust (highly reproducible), simple, and versatile indoor behavioral experimental systems other than the olfactory PER associative learning paradigm is important for the rise of honey bee behavioral genetics.

We focused on the antennal response (AR) of honey bees as one of these indoor behavioral experimental systems. Insect antennae are essential for various sensory receptions such as olfaction, gustation, and mechanoreception and are used to sense the external world (Hallem et al., 2005; Staudacher et al., 2005; Vogt and Riddiford, 1981). Various insect species move their antennae in response to odorants, visual stimuli, and mechanical stimuli (Honegger, 1981; Mamiya et al., 2011; Natesan et al., 2019; Staudacher et al., 2005). In honey bees, ARs to motion, odor, and mechanical stimuli and learning-dependent changes in AR to odors have been reported (Cholé et al., 2015; Erber and Kloppenburg, 1995; Erber et al., 1993; Gascue et al., 2022; Suzuki, 1975). In addition, antennal contact is essential in maintaining society through nestmate recognition and pheromone reception in eusocial insects, including honey bees (Carolina Gomez Ramirez et al., 2023; Ozaki et al., 2005; Sharma et al., 2015). To date, multi-animal tracking studies in eusocial insects using individual identification tags have revealed developmental changes in the social behaviors and responses of individuals to different social circumstances, which were inferred from individual positions inside the nest and inter-individual interactions (Ai et al., 2017; Crall et al., 2015; Crall et al., 2018; Liberti et al., 2022; Mersch et al., 2013; Stroeymeyt et al., 2018; Wario et al., 2015). However, by colony-level observations, determining which behavioral components change and affect individual responses is generally difficult. Therefore, developing a robust experimental system for the measurement and analysis of antennal movements in honey bees is expected to lead to a better understanding of the behavioral components that influence not only environmental recognition but also social behaviors.

In previous studies, the movements of insect antennae were measured by manually determining their position and angle for each frame of video or using phototransistors, which register the movement of the antennae (Erber and Kloppenburg, 1995; Erber et al., 1993; Okada and Toh, 2004). Recently, video analysis technologies have advanced, and methods for automatically tracking the movement of body parts of the honey bee have been reported, such as tracking the marked colors at the tip of the antennae or separating the outline of the antennae from the background by image processing(Cholé et al., 2015; Cholé et al., 2022; Roy Khurana and Sane, 2016). In addition, a markerless posture-tracking tool using deep learning, DeepLabCut (Mathis et al., 2018; Nath et al., 2019), has been widely used in various research fields, such as ecology and neuroscience (Mathis and Mathis, 2020), and is becoming recognized as a powerful tool that does not require special devices or expertise, in addition to being noninvasive and robust in analyzing videos with a complex background. Recently, DeepLabCut has also been used in honey bees, and the AR to odors with positive or negative valence was analyzed, revealing more accurate behavioral descriptions than traditional PER protocols which only output binary response (Gascue et al., 2022).

This study aimed to enhance the experimental system for analyzing the ARs of honey bees using DeepLabCut. We focused on the AR to motion stimuli reported in previous studies, in which bees tilted their antennae in the opposite direction to the upward and downward motion stimuli (Erber and Kloppenburg, 1995; Erber et al., 1993) and confirmed that this AR was successfully detected by tracking antennal movements using DeepLabCut. In addition, we revealed that ARs to forward and backward motion stimuli were observed in honey bees. An investigation of the developmental maturation of honey bee ARs showed that ARs to motion stimuli were not detectable in bees immediately after emergence but became mature through post-emergence development in an experience-independent manner. Furthermore, unsupervised clustering analysis using multidimensional data created by processing tracking data using DeepLabCut classified antennal movements into different clusters, suggesting the efficacy of data-driven analysis for the behavioral classification of honey bees.

## Materials and Methods

### Animal

European honey bee (*Apis mellifera* L.) colonies were purchased from Kumagaya Beekeeping Company (Saitama, Japan) and Okinawa Kariyushi apiary (Okinawa, Japan) and maintained at the University of Tokyo (Tokyo, Japan). Workers flying outside the hive were collected and used for experiments examining ARs to motion stimuli, and for the ‘flying (F)’ group in the experiment examining the developmental maturation of AR. To test the developmental maturation of AR, we collected dozens of newly emerged workers identified by their fuzzy appearance (Winston, 1987) from the hives and divided them into four groups; ‘newly emerged (NE)’ group whose AR was analyzed on the day of collection, ‘colony-reared (Cr)’ group which was returned to the colony without any treatments, ‘colony-reared with one wing removed (CrWR)’ group which had one wing removed and returned to the colony, and ‘incubator-reared (Ir)’ group which was maintained in an acryl cage (95 × 55 × 110 mm in size) with a piece of honeycomb and approximately 30 workers captured from the hive, and kept in an incubator at 34 °C with honey and water fed *ad libitum*. Bees in the different groups were marked on their thoraxes with different colors using paint markers (POSCA, Japan). After rearing in the hive or incubator for ten days, bees in the Cr, CrWR, and Ir groups were collected from the hive or acrylic cage and used in the experiment.

### Recording antennal movement during presenting motion stimuli to bees

The collected workers were anesthetized on ice and fixed in a P-1000 pipette tip with masking tape. After awakening from anesthesia, bees were fed 4 µL of 30% (w/v) sucrose solution. 6 mm wide black and white vertical stripes were displayed on two LED monitors (60 Hz refresh rate) placed facing each other at a 30° angle, and motion stimuli were presented to a bee set between the monitors by moving these stripes at 24 mm s^-1^ using PowerPoint animation (Fig. 1). Bees were set vertically when vertical motion stimuli were presented, or set horizontally when horizontal motion stimuli were presented. After familiarization for at least 3 min, each of the vertical or horizontal motion stimuli was presented alternately twice with an interstimulus interval of approximately 10 s (Fig. 1A). The heads of the bees were recorded at 30 fps (frames per second) with a resolution of 1024 × 768 by a Raspberry pi camera module (Kumantech) set above the bees during the experiment. To capture the entire antenna, the camera was tilted at 20° when the bees were set horizontally (Fig. 1D).

**Figure 1.**
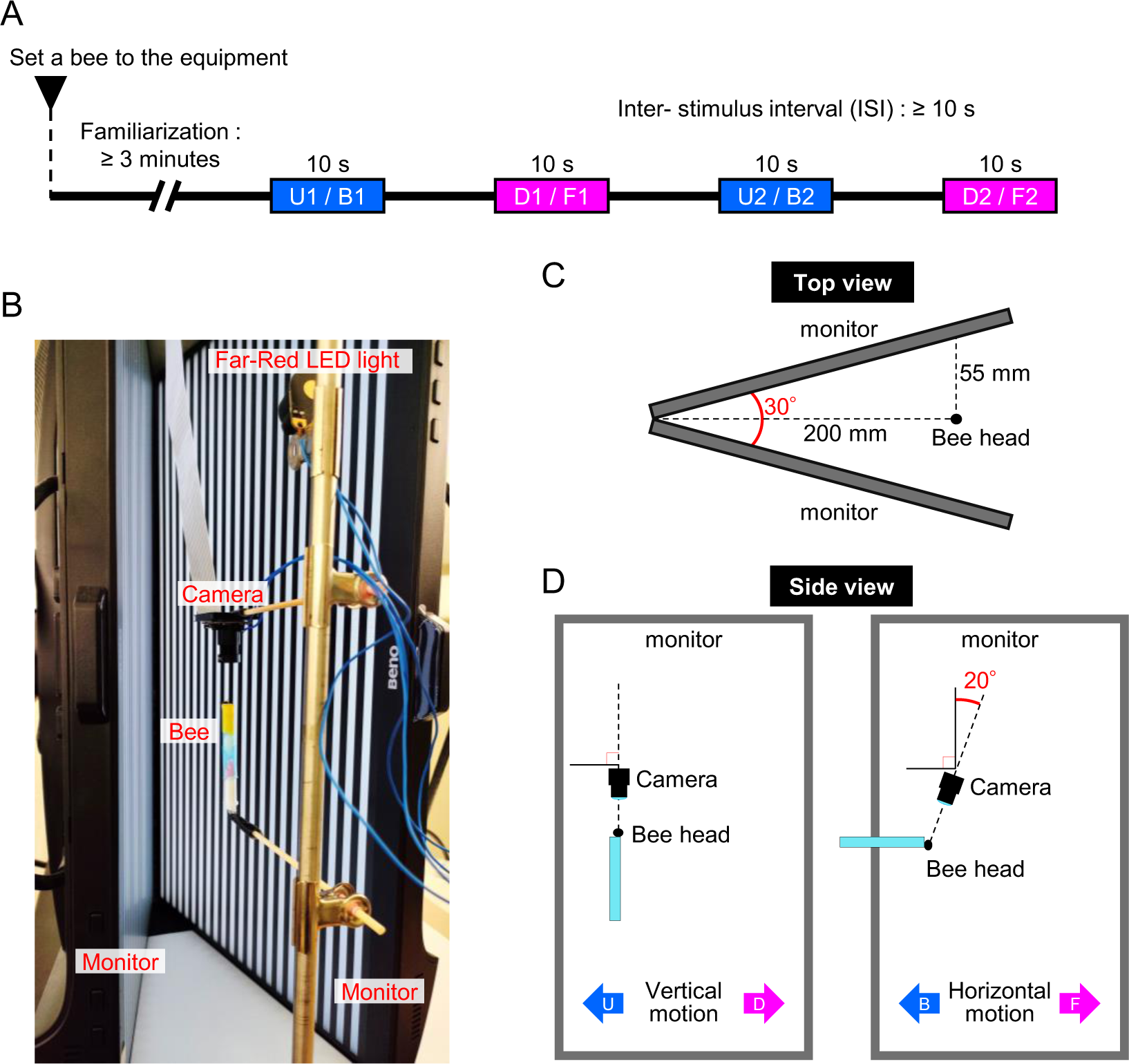
Overview of the experimental setup for the presentation of motion stimuli to a fixed bee. **(A)** The time course for the experiments. Blue boxes indicate the periods during the presentation of upward (U1 and U2) or backward (B1 and B2) motion stimuli, and magenta boxes indicate the periods during the presentation of downward (D1 and D2) or forward (U1 and U2) motion stimuli. **(B)** A picture of the experimental setup. **(C)** Schematic diagram of the experimental setup viewed from the top. The positions of monitors and a bee are shown. **(D)** Schematic diagrams of the side view of the experimental setup in the experiment for the presentation of vertical (left) or horizontal (right) motion.

### Automatic tracking of antennal movement using DeepLabCut

Antennal movements were tracked using the DeepLabCut software (DeepLabCut GUI v2.2.2)(Mathis et al., 2018; Nath et al., 2019). A total of 200 frames were extracted from videos of 10 individuals (20 frames per individual) to create a training dataset for each vertical or horizontal motion presentation experiment. The following eight points were manually labeled on each extracted frame: the base, middle (the joint between the scape and flagellum), and tip of each antenna, central ocellus, and ventral center (for the vertical motion presentation experiment), or posterior center (for the horizontal motion presentation experiment). We trained the networks using these data with default parameter settings (ResNet50, 500,000 iterations) and evaluated the trained networks by checking the labeled frames. Frames with low-likelihood labeling were extracted for additional training, and the networks were retrained after manually correcting the mislabeled points with the same settings as in the first training. The networks created were used to track the antennal movement in each video.

### Data analysis

Data analyses were performed using R version 4.1.0. R package ‘dlcpr’ was used to load the tracking data for each stimulus for each individual into tidy data. Tracked points with a low likelihood (<0.90) were linearly interpolated from the points in the previous and following frames using the ‘imputeTS’ package. The angle of each antenna was calculated as the angle of the line connecting the base and tip of each antenna from the centerline of the head connecting the central ocellus and the ventral center (for the vertical motion presentation experiment) or the posterior center (for the horizontal motion presentation experiment). Circular histograms of all the frames for each stimulus were drawn using the ‘circular’ package. One-way analysis of variance (ANOVA) and post hoc Tukey’s honest significant difference (HSD) tests were performed using the average antennal angle of both antennae of each individual during the presentation of each stimulus. Unsupervised clustering was performed using data obtained by tracking the full-length video of each individual in the vertical motion presentation experiment. To standardize each frame, all tracking points were translated such that the dorsal center was at the origin and rotated such that the ventral center overlapped with the y-axis, followed by scaling to make the distance from the dorsal center to the ventral center constant. For every 10 frames (0.33 s), the maximum, minimum, mean, and standard deviation of the x- and y-coordinates were calculated for eight points: the base, middle, and tip of each antenna and the centroid of these three points for each antenna. The resulting data points, with a total of 64 dimensions, were used to determine the appropriate number of clusters by calculating the gap statistic (Tibshirani et al., 2001) and were clustered into eight clusters using Hartigan–Wong’s k-means algorithm. Clustering results were visualized using a uniform manifold approximation and projection (UMAP). The corresponding stimulus (spontaneous, downward, or upward) in each frame was manually annotated.

## Results

### Antennal response to motion stimuli

First, we examined whether the ARs to upward and downward motion stimuli, reported in previous studies (Erber and Kloppenburg, 1995; Erber et al., 1993), could be detected even in our experimental system by automated tracking using DeepLabCut. Vertical motion stimuli were presented to a fixed worker between the monitors (Figs. 1C, D, and 2A). The movements of the left and right antennae were tracked using DeepLabCut during the presentation of each stimulus, and the angles of the antennae from the centerline of the head in the horizontal plane were calculated (Fig. 2B, Movie 1). The angular distribution of all frames of the antennae during the presentation of upward or downward motion stimuli showed a tendency to move the antennae in the opposite direction of the motion stimuli (Fig. 2C). The angular distribution of the spontaneous antennal movement was rather similar to that of the downward motion stimulus. However, the average angle of all frames of ‘spontaneous’ movement was in between the average angles during ‘upward’ and ‘downward’ motion presentation (Fig. 2C). Furthermore, we calculated the average angles of both the antennae of each bee for each stimulus and found significant differences in the angles between the stimuli: approximately 90°, 60°, and 100° for spontaneous movement, responses to upward and downward motion stimuli, respectively (Fig. 2D). These results are consistent with those of previous studies (Erber and Kloppenburg, 1995; Erber et al., 1993), confirming the applicability of our automated tracking system.

**Figure 2.**
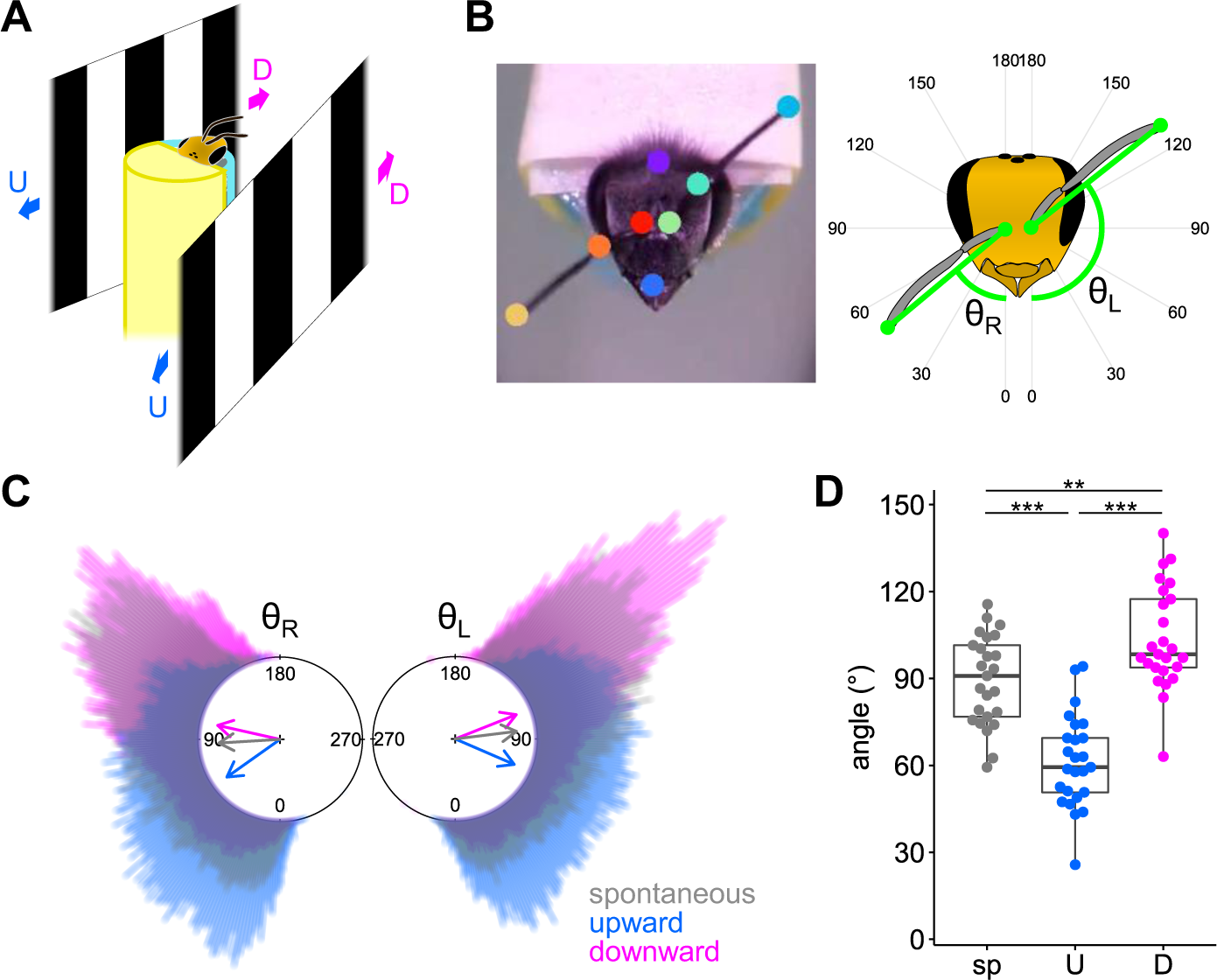
ARs to vertical motion stimuli. **(A)** Schematic diagram of the experimental setup to present the vertical motion stimuli displayed on the monitors to a fixed honey bee. Magenta and blue arrows indicate downward and upward motions, respectively. **(B)** One frame of the video after processing using DeepLabCut (left) and the schematic diagram of angles of the left (θ_L_) and right (θ_R_) antennae (right). **(C)** Circular histogram of θ_L_ andθ_R_ for all frames (15,000 frames per stimulus; 25 individuals, 10 sec × 2 for each stimulus, recording at 30 fps) of spontaneous movements or during the presentation of the upward or downward motion stimulus. The angles of arrows in the circular histograms indicate the average antennal angles of all frames for each stimulus, and their lengths indicate the lengths of the summed unit vectors of the antennal angles for all frames divided by the number of frames. **(D)** Comparison of ARs among stimuli. Each dot represents the average angle of both antennae for each individual while each motion stimulus was presented. n = 25 for each stimulus. ***: p < 0.001, **: p < 0.01 by Tukey’s HSD test.

Because the previous study only examined the ARs to vertical motion stimuli by measuring the antennal angles in the transverse plane, we next examined whether the AR was also observed in response to backward and forward motion stimuli by measuring the antennal angles in the coronal plane. A worker fixed in a plastic tip was placed horizontally between the monitors to present horizontal motion stimuli (Figs. 1C, D, and 3A). Antennal movements were tracked as in the experiment for AR to vertical motion stimuli, and the angles of the antennae from the centerline in the coronal plane were calculated (Fig. 3B, Movie 2). As in the AR to vertical motion stimuli, the angular histogram of all video frames during the presentation of each stimulus showed a tendency to move the antennae in the opposite direction to the motion stimuli, and the average angles of the antennae spontaneously moving were between those during the presentation of each stimulus (Fig. 3C). The average angles of both antennae of each bee also differed significantly between the stimuli: approximately 65°, 45°, and 70° for spontaneous movement, responses to backward and forward motion stimuli, respectively (Fig. 3D). The difference in the average angles for different stimuli in the coronal plane was smaller than that in the horizontal plane because of the narrower range of antennal movements in the coronal plane. These results indicate that workers respond to external motion stimuli by moving their antennae in the direction opposite to the motion in at least four directions: upward, downward, backward, and forward.

**Figure 3.**
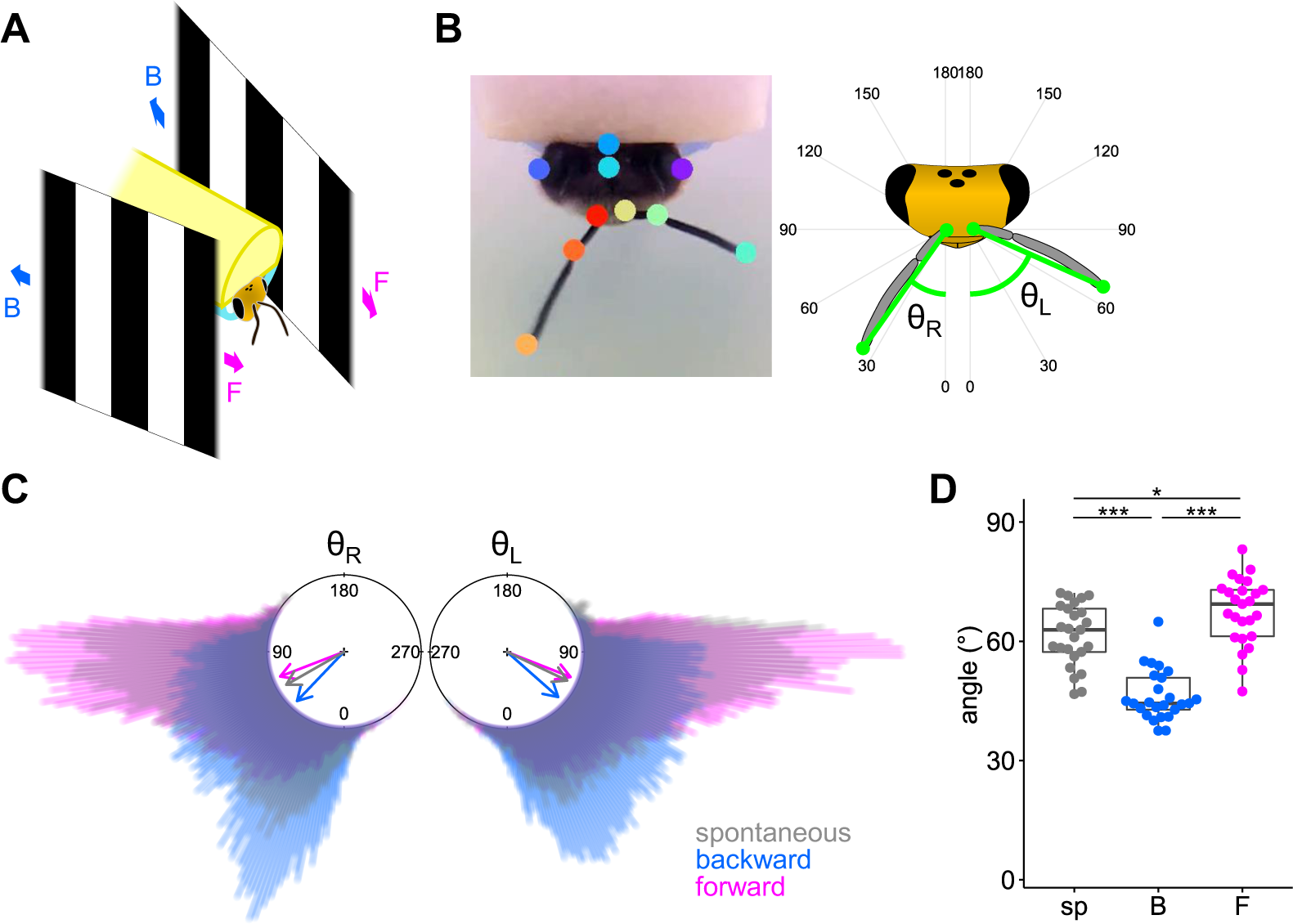
ARs to horizontal motion stimuli. **(A)** Schematic diagram of the experimental setup to present the horizontal motion stimuli displayed on the monitors to a fixed honey bee. Magenta and blue arrows indicate forward and backward motions, respectively. **(B)** One frame of the video after processing using DeepLabCut (left) and the schematic diagram of angles of the left (θ_L_) and right (θ_R_) antennae (right). **(C)** Circular histogram of θ_L_ and θ_R_ for all frames (15,000 frames per stimulus; 25 individuals, 10 sec × 2 for each stimulus, recording at 30 fps) of spontaneous movements or during the presentation of the backward or forward motion stimulus. The angles of arrows in the circular histograms indicate the average antennal angles of all frames for each stimulus, and their lengths indicate the lengths of the summed unit vectors of the antennal angles for all frames divided by the number of frames. **(D)** Comparison of antennal responses among stimuli. Each dot represents the average angle of both antennae for each individual while each motion stimulus was presented. n = 25 for each stimulus. ***: p < 0.001, *: p < 0.05 by Tukey’s HSD test.

### Developmental maturation of AR after emergence

A previous study reported that ARs to odorants were limited in newly emerged workers and acquired in the adult life (Cholé et al., 2022). Thus, we next compared the AR to the vertical motion stimuli of newly merged (NE) and flying (F) workers to investigate whether ARs to visual stimuli also mature after emergence. We calculated the direction-specific antennal response (DAR), which is the angular difference between the average angle of the antennae during upward and downward motion stimuli presentation for each bee (Fig. 4A)(Erber and Kloppenburg, 1995; Erber et al., 1993), and compared the results between groups. The results showed that F had a significantly larger DAR than NE, which had almost zero DAR (Fig. 4B). The NE showed similar angular distributions for all frames and average angles during the presentation of each stimulus (Fig. S1A, B), which was contrary to the results obtained for F (Fig. S1C, D).

**Figure 4.**
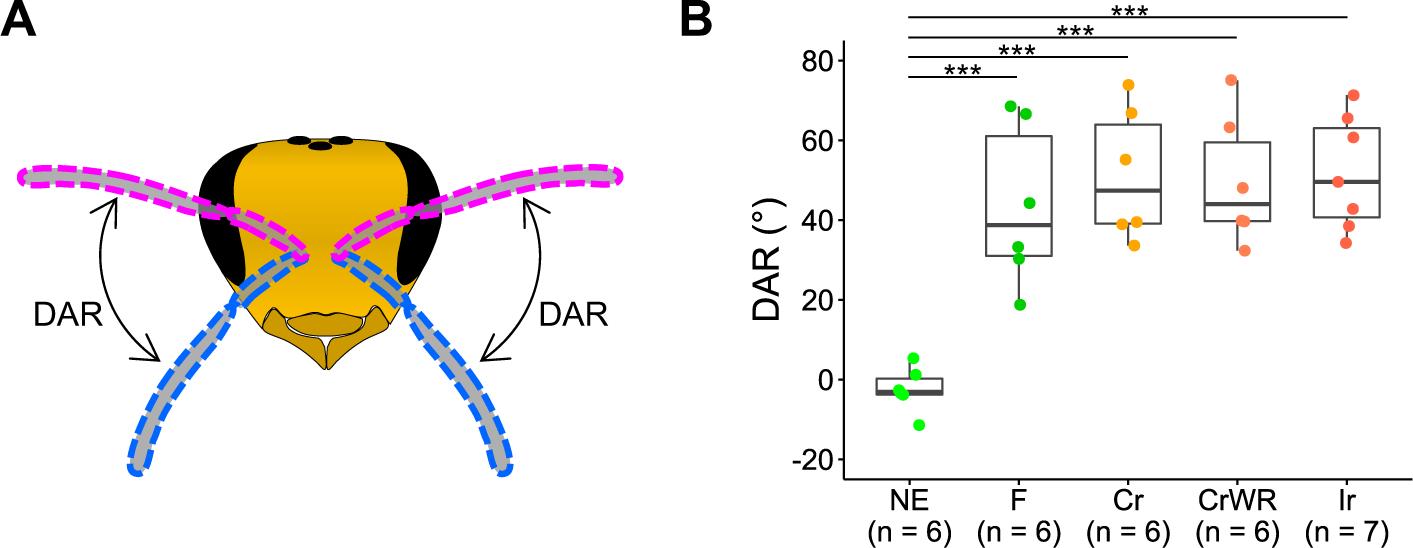
Development of AR to vertical motion stimuli. **(A)** Schematic diagram of DAR. The difference between the mean angles of antennae during the presentation of upward and downward motions is defined as DAR. **(B)** Comparison of DARs of bees in the groups analyzed at different developmental stages (NE, F) or with different treatments (Cr, CrWR, Ir). ***: p < 0.001 by Tukey’s HSD test.

As the motion presented to the compound eye is opposite to the direction in which a bee moves under natural conditions, moving the antennae in the direction opposite to the motion stimuli displayed on the monitors in this experimental system implies that bees were trying to move their antennae to perceive sensory information in the direction in which they are moving, for example, to recognize objects around the landing site (Evangelista et al., 2010). Therefore, the difference in DAR between the NE and F groups suggests that AR to motion stimuli is acquired based on each individual’s flight experience or developmental maturation after emergence. To test these possibilities, we divided newly emerged workers collected from a hive into three groups and reared them under different conditions: 1) those that were returned to the same hive from which they were collected with no treatment (colony-reared [Cr]), 2) those that were returned to the same hive from which they were collected with two wings on one side removed (colony-reared with wings removed [CrWR]), and 3) those that were reared in a small acrylic cage containing approximately 30 adult workers in an incubator (incubator-reared [Ir]). The bees in the CrWR and Ir groups could not fly, thus, they could not acquire AR through experience. The ARs to motion stimuli of bees in the Cr, CrWR, and Ir groups were analyzed 10 days after the treatments because the first orientating flight of workers was reported to occur as early as 4 days old (Winston, 1987), and bees in the Cr group were considered to have flight experience. The results showed that the DARs of bees in the Cr, CrWR, and Ir groups were significantly different from those in the NE group. In contrast, they were comparable to that of the F group and not significantly different from each other (Fig. 4B). In addition, the average angle of the antennae during the presentation of each stimulus was not constant across stimuli in the Cr, CrWR, and Ir groups (Fig. S1E-J). These results indicate that AR to motion develops with age in a flight experience-independent manner, suggesting that this response is an innate honey bee behavior.

### Classification of AR by unsupervised clustering

Animal tracking data can be used for unsupervised behavioral classification (Fujimori et al., 2020; Huang et al., 2021; Segalin et al., 2021). Therefore, we examined whether data-driven analysis could classify the tracking data of the antennal movements. Antennal movements were tracked using DeepLabCut for all video frames of 25 individuals in the vertical motion presentation experiment. After the tracking points of each frame were standardized (see Methods for details), the maximum, minimum, mean, and standard deviation of the x- and y-coordinates were calculated every 10 frames (0.33 s) for the following eight points: the base, middle, and tip of each antenna and the centroid of these three points. The resulting data points with 64 dimensions were divided into eight clusters and visualized using UMAP (Fig. 5A). While data points for spontaneous movements were distributed almost uniformly on the UMAP, those during the presentation of upward and downward motion stimuli were distributed nonuniformly, with gradients in opposite directions (Fig. 5B). The proportions of data points for spontaneous movements included in each cluster were almost uniform (approximately 50% in each cluster), whereas those during the presentation of the downward and upward motion stimuli varied across clusters (Fig. 5C). As each video contained frames from over 10 s before the first stimulus presentation to a few seconds after the last stimulus, the number of data points corresponding to spontaneous movements was greater than the sum of the number of data points during the presentation of the upward and downward motion stimuli. In addition, we examined how the data points corresponding to spontaneous movements or during the presentation of upward and downward motion stimuli were distributed into eight clusters. The data points for spontaneous movements were somewhat uniformly classified into all clusters, with a slight tendency to be more classified into Cluster 3 and less classified into Cluster 2 (Fig. 5D). The data points during the presentation of the upward motion stimulus were prominently classified into Clusters 2 and 4 and less frequently classified into Cluster 3 (Fig. 5E). The data points during the presentation of the downward motion stimulus were more classified into Cluster 3 and less into Clusters 2 and 3, showing the opposite tendency to that during the presentation of the upward motion stimulus (Fig. 5F). All clusters included data points from at least 23 of the 25 individuals, and there were no clusters with a high percentage of data points from any particular individual, suggesting that the biases described above were not due to variations among individuals, such as variations in the AR, the angle at which bees were fixed in the experimental setup, or the distance from the head of the bee to the monitors (Fig. S2).

**Figure 5.**
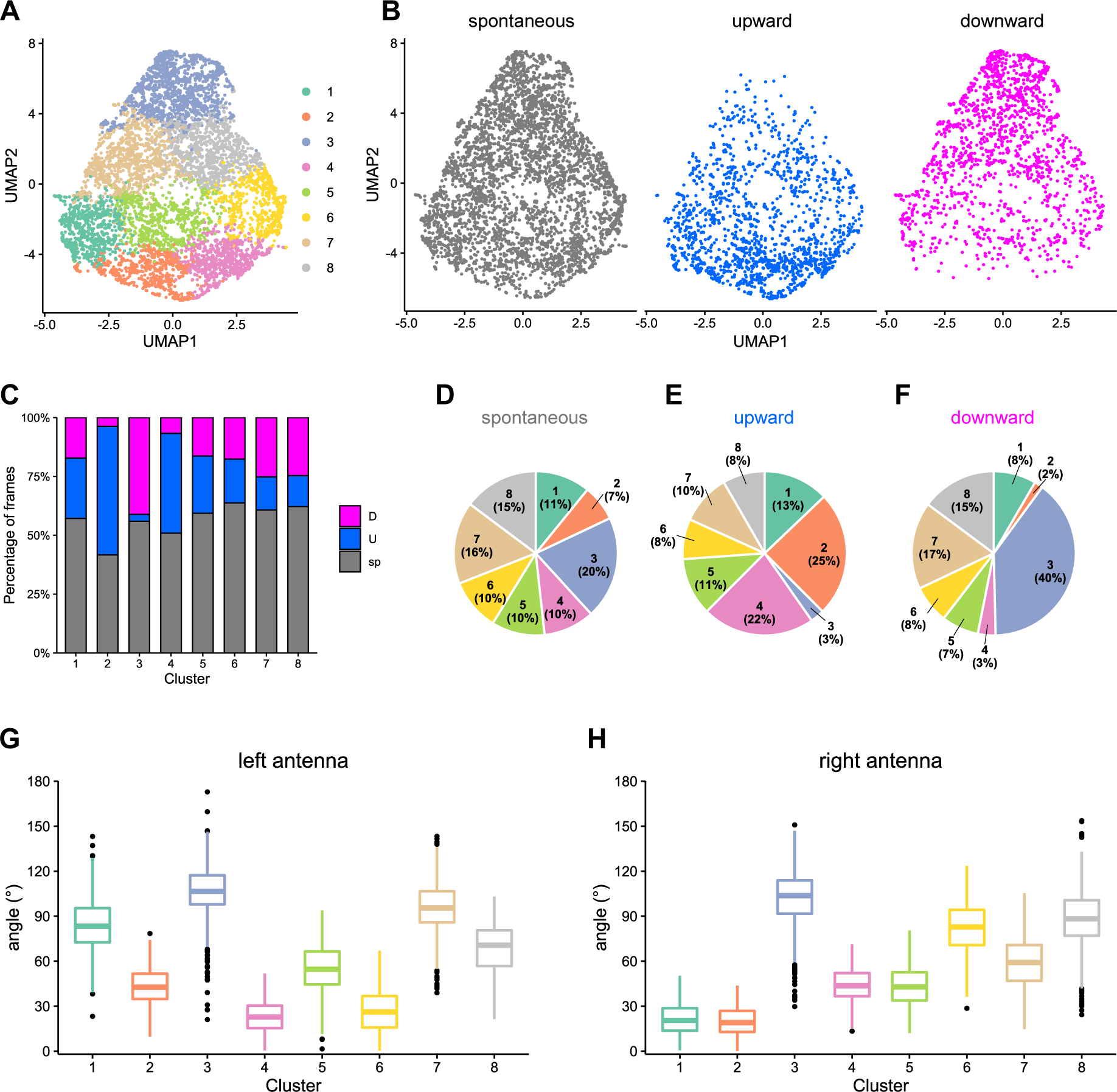
Unsupervised clustering of antennal movements. **(A)** UMAP visualization of clusters of data points processed from tracking data of all individuals for every 10 frames. **(B)** Distribution of data points corresponding to spontaneous movements (gray) or during the presentation of upward motion (blue) and downward motion (magenta) on the UMAP. **(C)** The proportion of data points corresponding to each stimulus in each cluster. Gray indicates the proportion of spontaneous movements, whereas blue and magenta indicate that of downward and upward motion stimuli, respectively. **(D-F)** Pie charts of the proportions of data points classified into each cluster among all data points corresponding to spontaneous movements (D), upward motion stimulus (E), and downward motion stimulus (F). **(G, H)** Boxplots of the left (G) and right (H) antennae angles for each cluster. The antennal angles of each data point were calculated from the averages of the x- and y-coordinates of the tip of each antenna.

Since the distribution of data points across clusters and the average antennal angle of each individual varied depending on the presented motion stimuli (Figs. 2C, D, and 5B-C), we measured the antennal angle for each cluster to examine the properties that characterize each cluster. Because we noticed that the bees often moved their left and right antennae separately when observing the frames corresponding to each cluster, we calculated the angle of each antenna. The antennal angles varied among clusters; for example, the angles of both antennae in Cluster 3 tended to be large, whereas those in Clusters 2 and 4 were small, and the difference between the angles of the left and right antennae in Clusters 1 and 6 was large compared to the other clusters (Fig. 5G, H). Reasonably, the data points in Cluster 3 had large angles for both antennae, and those in Clusters 2 and 4 had small angles, considering the differences in the distribution of data points (Fig. 5C, E, F) and the average angles (Fig. 2D) during the presentation of upward and downward motion stimuli. These results suggest that a data-driven analysis using data points processed from tracking data can be used to classify antennal movements based on their characteristics.

## Discussion

We investigated AR to motion stimuli in a honey bee using a video analysis software based on deep learning, DeepLabCut. Our video analysis also successfully detected ARs in honey bees; honey bees moved their antennae in the direction opposite to the vertical motion stimuli, as reported in previous studies (Erber and Kloppenburg, 1995; Erber et al., 1993). In addition, honey bees can exhibit AR to horizontal motion stimuli by moving their antennae in the direction opposite to the motion stimuli in the coronal plane. However, compared with previous studies (Erber and Kloppenburg, 1995; Erber et al., 1993), the antennal angles and DAR, when presented with upward and downward motion stimuli, tended to be larger in the present study. This might be due to differences in the experimental setup, such as the width of the stripes used to present motion stimuli, the distance between the monitors and bees, and the speed of the motion stimuli. In addition, this could be due to the difference in the measurement devices because previous studies used only four phototransistors to measure the angles of each antenna and could not measure angles with as fine a resolution as the motion tracking by DeepLabCut in this study. However, the tendency to move the antennae in the direction opposite to the motion stimulus is consistent between previous and present studies, showing that AR to motion is a robust behavioral property.

AR to motion has been observed in flying insect species other than the honey bee, such as fruit flies and hawk moths, and is believed to play an important role in sensory perception during flight, in combination with AR to other sensory information (Krishnan and Sane, 2014; Mamiya et al., 2011). Although we examined AR only to motion at a constant speed (visual stimulus) in the present study, previous studies have examined how insects control their antennal position during exposure to airflow (physical stimuli) or airflow and visual stimuli together (Natesan et al., 2019; Roy Khurana and Sane, 2016). Considering that a recent study used DeepLabCut to examine AR to odors with positive or negative valence in honey bees (Gascue et al., 2022), the experimental setup coupled with automated tracking using DeepLabCut used in this study could easily be expanded to examine AR to multimodal sensory information by simultaneously presenting motion (visual stimulus), airflow (physical stimulus), and odor (olfactory stimulus). This will deepen our understanding of how insects control their antennal movements to properly acquire sensory feedback in nature.

In honey bees, direction-specific AR to motion stimuli has been reported to be affected by the administration of serotonin and octopamine in the optic lobes, which are the primary visual centers in the insect brain. The effect of serotonin on AR was observed when injected into the lobula, medulla, and lamina, whereas that of octopamine was observed only when injected into the lobula, suggesting a particularly important role of the lobula for AR in motion stimuli (Erber and Kloppenburg, 1995). In *Drosophila*, some lobula neurons respond specifically in each of four directions (upward, downward, forward, and backward) (Fischbach and Dittrich, 1989). Notably, the motor neurons that control antennal movement in the honey bee extend dendrites to the dorsal lobe of the antennal lobe, where projections from the lobula terminate (Kloppenburg, 1995; Maronde, 1991). This study revealed the presence of AR to each of the four motion directions (upward, downward, forward, and backward), suggesting that AR may arise in response to direction-specific neuronal input from the lobula to the motor neurons regulating antennal movements. This study also revealed that ARs to motion stimuli are not experience-dependent but are acquired through individual development after emergence. Because workers engage in in-hive tasks for a certain period after emergence (Seeley, 1995; Winston, 1987) and do not need to respond to visual stimuli, the inability of newly emerged workers to respond to motion stimuli is not expected to significantly impact their survival rate. Since previous immunohistochemical studies have revealed the presence of serotonin, serotonin receptor 5-HT_1A_, and octopamine in the lobula of the honey bee (Schürmann and Klemm, 1984; Sinakevitch et al., 2005; Thamm et al., 2010), investigating changes in their expression patterns after emergence in correlation with the development of AR can contribute to elucidating the molecular and neural mechanisms of AR to motion stimuli.

Finally, the data points processed from the tracking data using DeepLabCut were classified into clusters with different antennal movement characteristics by data-driven analysis. Contrary to the analysis focusing on the antennal angle in each frame, in which the responses of the left and right antennae were averaged and the differences between them could not be detected (Fig. 2), data-driven analysis successfully classified the data points with different left and right antennal movements into different clusters, indicating the efficacy of data-driven analysis without relying on any hypotheses. Honey bees use their antennae not only to perceive sensory cues from the environment but also to interact with nestmates (Winston, 1987). Therefore, tracking antennal movements during inter-individual interactions and subsequent data-driven clustering may contribute to unveiling the details of how nest-mate recognition and interactions between individuals engaging in different tasks are reflected in antennal movements. Recently, behavior classification by data-driven analysis has been used to explore behavior-related brain regions in combination with neural activity mapping, or to analyze the phenotypes of individuals whose gene functions and neuronal activities are manipulated (Huang et al., 2021; Markowitz et al., 2023). Since genome editing and transgenesis have recently been utilized in the honey bee (Carcaud et al., 2023; Chen et al., 2021; Cheng et al., 2023; Değirmenci et al., 2020; Kohno and Kubo, 2018; Nie et al., 2021; Roth et al., 2019; Wang et al., 2021), it is expected that the data-driven behavioral analysis could contribute to the analysis of the molecular and neural basis of social behaviors of the honey bee, which are still largely unknown, in the near future.

## Competing interests

The authors declare no competing interests.

## Funding

This work was supported by Japan Society for the Promotion of Science (JSPS) KAKENHI Grant Number 20K15839 (Grant-in-Aid for Early-Career Scientists).

## Notes

### Competing Interest Statement

The authors have declared no competing interest.

